# Centromere Detection of Human Metaphase Chromosome Images using a Candidate Based Method

**DOI:** 10.1101/032110

**Authors:** Akila Subasinghe, Jagath Samarabandu, Yanxin Li, Ruth Wilkins, Farrah Flegal, Joan H. Knoll, Peter K. Rogan

## Abstract

Accurate detection of the human metaphase chromosome centromere is an critical element of cytogenetic diagnostic techniques, including chromosome enumeration, karyotyping and radiation biodosimetry. Existing image processing methods can perform poorly in the presence of irregular boundaries, shape variations and premature sister chromatid separation, which can adversely affect centromere localization. We present a centromere detection algorithm that uses a novel profile thickness measurement technique on irregular chromosome structures defined by contour partitioning. Our algorithm generates a set of centromere candidates which are then evaluated based on a set of features derived from images of chromosomes. Our method also partitions the chromosome contour to isolate its telomere regions and then detects and corrects for sister chromatid separation. When tested with a chromosome database consisting of 1400 chromosomes collected from 40 metaphase cell images, the candidate based centromere detection algorithm was able to accurately localize 1220 centromere locations yielding a detection accuracy of 87%. We also introduce a Candidate Based Centromere Confidence (CBCC) metric which indicates an approximate confidence value of a given centromere detection and can be readily extended into other candidate related detection problems.

## 1 Introduction

The centromere of a human chromosome (figure 1) is the primary constriction to which the spindle fiber is attached during the cell division cycle (mitosis). The detection of this salient point is the key to calculating the centromere index which can lead to the type and the number of a given chromosome. The reliable detection of the centromere by image analysis techniques is challenging due to the high morphological variations of chromosomes on microscope slides. This variation is caused by various cell preparatory and staining methods along with many other factors during mitosis. Irregular boundaries and large variations in morphology of the chromosome can cause a detection algorithm to miss the constriction, especially in higher resolution chromosomes. Premature sister chromatid separation can also pose a significant challenge, since the degree of separation varies from cell to cell, and even among chromosomes in the same cell. In such cases, the width constriction can be missed by image processing algorithms, and can result in incorrect localization of a centromere on one of the sister chromatids.

**Fig. 1.**
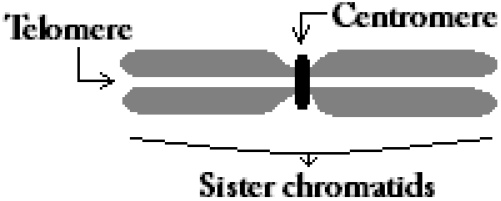
Demonstrates the anatomy of a human metaphase chromosome using a simple graphical design with key components labeled.

From an image analysis perspective, the high morphological variations in human chromosomes due to their non rigid nature pose a significant challenge. Cell preparation and staining techniques and also vary among on the laboratories. The end results obtained from clinical cytogenetic vs. reference biodosimetry laboratories can produce chromosome images that differ significantly in their appearance. As an example, chromosomes that were DAPI (4’,6-Diamidino-2-Phenylindole) stained would demonstrate different intensity features and boundary characteristics from chromosomes subjected only to Gei-msa staining. Additionally, the stage of metaphase in which the cells were arrested along with environmental factors such as humidity can dictate the shape characteristics of each individual cell and introduce a large variance to the data set. Furthermore, in some preparatory methods, the cells are denatured introduce significant noise at the chromosome boundary. These same factors can also dictate the amount of premature sister chromatid separation in some of the cells. Effective algorithms for centromere detection need to be able to handle the high degree of shape variability present in different chromosomes, while correcting for artifacts such as premature sister chromatid separation. Figure 2 below illustrates a sample set of shapes of chromosomes in the data set and their high morphological variations.

**Fig. 2.**
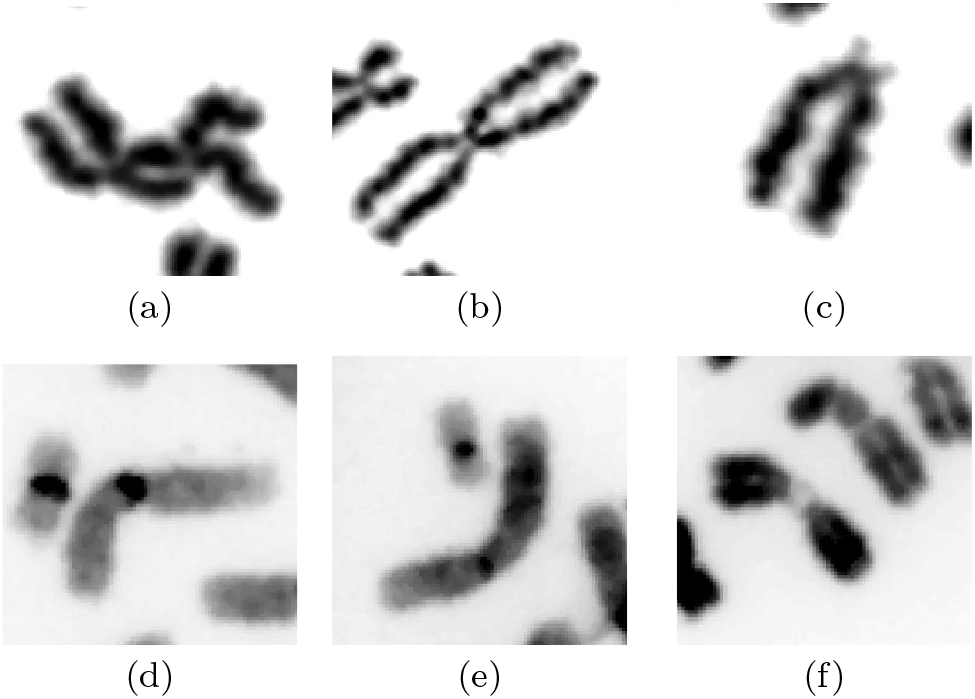
Depicts various degrees of sister chromatid separation present in some Geimsa stained chromosome cell images (fig 2(a), (b) & (c)) as well as some lengthy chromosomes characteristic to those prepared at a cytogenetic laboratory (fig 2(d), (e) & (f)).

This research is a prerequisite for the development of a set of algorithms for detecting dicentric chromosomes (possessing two centromeres) which are diagnostic of radiation exposures in cytogenetic biodosimetry. The ability of the proposed algorithm to handle high degrees of morphological variations and also to detect and correct for the artifact created by premature sister chromatid separation in cell images is also critical to detecting dicentric chromosomal abnormalities.

Numerous computer algorithms have been proposed over time for chromosome analysis ranging from metaphase finding [1], Karyotype analysis [2] to centromere and dicentric detection [3], [4]. These methods are either constrained by the protocol used for staining the cell image or by the morphology of the chromosome. We have previously proposed an algorithm to locate the centromere by calculating a centerline with no spurious branches irrespective of boundary irregularities or the morphology of the chromosome [5]. Mohammad proposed an approach where he used our previous approach to derive the centerline and then used a curvature measure to localize the centromere location instead of the width measurements [6]. Another interesting approach by Jahani and Setarehdan involves artificially straightening chromosomes prior to creating the trellis structure using the centerline derived through morphological thinning [7]. Yet all these methods, including our previous approach, work well only with smooth object boundaries. The absence of a smooth boundary will directly affect the centerline and thus make the feature calculations noisy. Furthermore, the accuracy of all these methods is adversely impacted by sister chromatid separation.

We propose a candidate based centromere localization algorithm capable of processing highly bent chromosome images prepared with a variety of staining techniques. This method can also detect and correct for the artifacts introduced by premature sister chromatid separation. Since centerline-based methods tend to perform better than other methods, we have proposed an algorithm which utilizes the centerline simply to divide the chromosome contour into two nearly symmetric partitions, rather than using this feature as a basis for width measurements. In centerline-based methods, boundary irregularities often get embedded in the centerline, which therefore introduces noise into the width profiles. By avoiding the centerline as the basis for measurements, the condition of the chromosome boundary does not impact the smoothness of width profile measurements. Then, a Laplacian-based algorithm that integrates intensity measurements in a weighting scheme, biases the thickness measurement by tracing vectors across regions of homogeneous intensity. To address image processing artifacts arising from sister chromosome separation, an improved contour partitioning algorithm is presented. This paper also introduces the Candidate-Based Centromere Confidence (CBCC) metric, which measures the confidence in an accurate centromere assignment [8]. This metric is used in tests of the algorithm on a large data set of chromosomes, with the aim of validating the performance of the algorithm.

## 2 Methods

The following section will describe the proposed candidate based centromere detection algorithm in detail. This method can be functionally divided into following sections for clarity,

– Segmentation & centerline extraction
– Contour partitioning & correcting for sister chromatid separation
– Chromosome thickness measurement using the intensity integrated Laplacian method
– Candidate point generation & metaphase centromere detection

The proposed method operates on well-separated chromosomes that do not overlap or touch others. To ensure that the metaphase cells with the maximum number of segmentable chromosomes are analyzed, cells with incomplete chromosome complements and those with higher densities of overlapping and touching chromosomes are initially deprecated using a content-based classification procedure [9].

To develop the method, individual chromosomes were first selected manually, while the remainder of the process has been automated. Our Automated Dicentric Chromosome Identifier (ADCI) software also automatically selects individual chromosomes [10]. Once a chromosome is selected, the proposed method segments this object by local thresholding [11] and Gradient Vector Flow (GVF) active contours [12]. Upon extraction based on the GVF boundaries, the contour of the chromosome is partitioned using a polygonal shape simplification algorithm known as Discrete Curve Evolution (DCE) [13] which simplifies the shape of the object by iteratively deleting vertices based on their importance to the overall shape of the object. Then a Support Vector Machine (SVM) classifier is used to pick the best set of points to isolate the telomeric regions i.e. at the ends of the chromosome. The segmented telomere regions are then tested for evidence of sister chromatid separation using a second trained SVM classifier designed to capture shape characteristics of the telomere regions and then corrected for that artifact. Afterwards, the chromosome is split into two partitions along the axis of symmetry and a modified Laplacian based thickness measurement algorithm (called Intensity Integrated Laplacian or IIL) is used to calculate the width profile of the chromosome [8]. This profile is then used to identify a possible set of candidates for centromere location(s) and features are calculated for each of those locations. Next, a third classifier is trained on expert-classified chromosomes to detect centromere locations in test chromosomes. In most instances, each chromosome will contain at least one centromere. therefore, the correct centromere should be present among the candidates. The distance from the separating hyperplane is used as an indicator for the goodness of fit of a given candidate and thereafter used to select the best candidate from the pool of candidates. We test the hypothesis that best candidate for a given chromosome is a centromere location. In the following sections, we present further details of the algorithm. Section 2.1 and 2.3 have been previously published [8] and have been included in this paper to improve readability.

### 2.1 Segmentation & Centerline extraction

Once the user selects a point contained within a chromosome, a region of interest (ROI) containing the chromosome is selected and extracted. These regions will be processed separately in subsequent steps. Pre-processing consists of the application of a median filter followed by intensity normalization for the extracted window around the chromosome. The chromosome is first thresholded using Otsu's method and then the contour of that binary object is used as the initial contour for GVF active contours. The use of GVF active contours provides a contour that is smooth and that converges to boundary concavities [14].

In order to calculate the width profile of the chromosome using the thickness measuring algorithm, the chromosome contour is divided longitudinally into two approximately symmetric segments. We used Discrete Curve Evolution (DCE)-based skeletal pruning [15], [5] to obtain an accurate centerline. DCE is a polygon evolution algorithm which evolves through vertex deletion based on a relevance measurement [15]. The objective of this stage of the algorithm is to obtain the set of anchor points (end points of the centerline). This set of anchor points is denoted by *E^P^* (|*E^P^* | = 2), where |.| is the cardinality operator).

Since the minimum polygon is a triangle, the end result of this process is a skeleton traversing the length of the chromosome that is connected to a single spurious branch (3 branches). Throughout this paper, we use the supercript *P* to refer to various point sets on the chromosome object contour. If C ∊ ℝ^2^ is the contour of the chromosome, the DCE initial anchor points (skeletal end points) for the centerline are denoted by 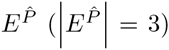. In order to obtain the centerline, the shortest branch of the resulting skeleton is pruned out to reveal the centerline of the chromosome. This yields the set of anchor points *E^P^*. This set of points is used for contour partitioning in the next section. The centerline was then shortened by 10% to discard any skeletal bifurcations that occur close to the end of the chromosome.

### 2.2 Contour partitioning & correcting for sister chromatid separation

Sister chromatid separation in chromosomes is an integral process that occurs during the metaphase stage of mitosis. Depending on the stage of mitosis at which the cells were arrested, varying degrees of sister chromatid separation may be evident. Furthermore colcemid, a chemical agent which is used mainly as a preparatory chemical in biodosimetry studies, can cause or exacerbate this condition and prematurely produce sister chromatid separation on metaphase cells. It is important that the algorithm and associated software be able to analyze chromosomes with sister chromatid separation.

In order to identify and correct for sister chromatid separation, we proposed an automated contour partitioning and shape matching algorithm. Our chromosome thickness measurement algorithm requires an approximate symmetric division of the contour of the chromosome. Accurate partitioning of the telomere region is necessary to identify evidence of sister chromatid separation and therefore correct for any such artifact as well as to split the contour into two segments accurately. Curvature of the contour is one of the most commonly used features for detecting salient points that can be used for partitioning [16]. An important requirement is that the location of these salient points needs to be highly repeatable under varying levels of object boundary noise. The DCE method described in the previous section was adapted to provide a set of initial salient points on the contour of the chromosome outline. This is because this method performs well with boundaries regardless whether they are smooth or not, yielding repeatable results [17]. The ability to terminate the process of DCE shape evolution at a given number of vertices further lends to its applicability. It was empirically established that a termination at 6 DCE points would ensure that the required telomere end points will be retained within the set of candidate salient points. Two of those 6 points will be selected as the end points of the centerline calculated in section 2.1. Contour partitioning is performed by selecting the best 4 point combination (including the two anchor points) that represents all the telomere end points.

The approach for selecting the optimal contour partitioning point combination occurs in two stages. Initially, a SVM classifier is trained to detect and label preferred combinations from the given 12 possible combinations for each chromosome. At this stage, all the combinations across the data set are used as a pool of candidates to train the classifier. Then, the signed Euclidian distance from the separating hyperplane (say ρ) is computed for each of the candidates for a given chromosome, considering only the combinations of that chromosome. This process ranks all the candidates according to the likelihood they are a preferred candidate. Unlike traditional rule-based ranking algorithms, this approach requires very little high level knowledge of the desirable characteristics. The positioning of the separating hyperplane encapsulates this high level information through user-specified ground truth. The highest-ranked candidate is the best combination of contour partitions for the given chromosome. The formal description of this procedure follows.

Features used for contour segmentation will be represented by *F^s^*, while a second set of features used for centromere localization will be denoted by *F^c^* (discussed in section 2.4).

Let *Φ_h_* be the curvature value at candidate point *h* and *S* ∊ ℝ^2^ be the skeleton of the chromosome with 6 DCE point stop criteria. We now define the following set of points,

– *D^P^* (⊂ C) is the set of six DCE vertices.
– *S^P^* constitutes of all the points in *D^P^* except the anchor points (*E^P^*). These are the four telomere endpoint candidates.

Then the family of sets *T^P^* for all possible combinations with the sets *E^P^* and *S^P^* would contain

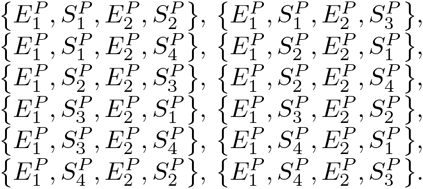

Figure 3 illustrates one such combination where the selected (connected by the blue line segments) combination for the contour partitioning points are given by 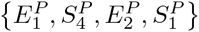.

**Fig. 3.**
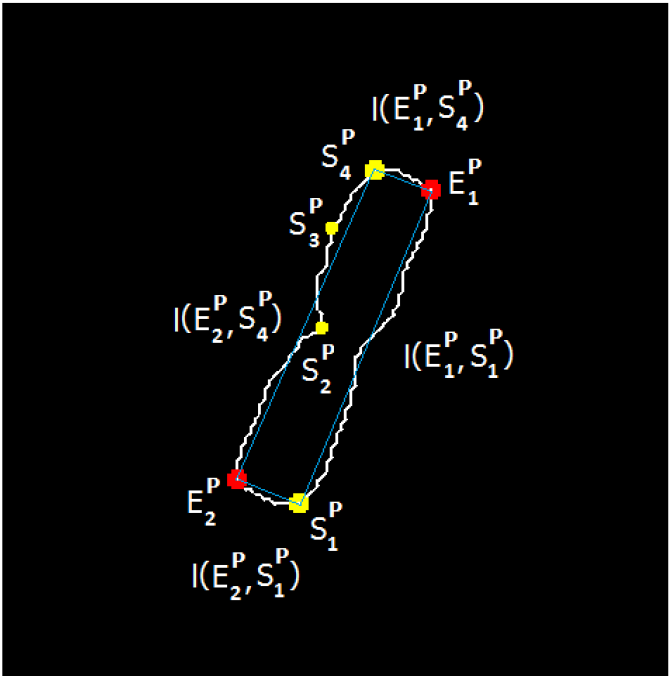
Demonstrates one possible combination for contour partitioning where the anchor point (red ‘+’ sign) 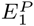 is connected with the candidate point 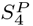 while the other anchor point 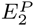 is connected with candidate point 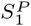 which captures the telomere regions. The (blue) line connects the set of points constituting the considered combination in this instance.

In order to identify the best possible combination for contour partitioning, we have used a SVM classifier trained with the 11 different features (*F^s^*) indicated below. Features 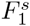 and 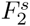 provide an indication to the saliency of the candidate point with respect to the skeletonization process. Features 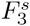 to 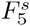 are three normalized features which capture the positioning of each candidate in the given combination. 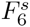 and 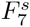 represent the shape or the morphology of the chromosome of interest (same values for all 12 combinations). The rationale behind the inclusion of these features is that they account for morphological variations across the cell images in the data set. 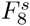 and 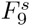 represent the curvature of the candidate points as well as the concavity/convexity of those locations. The features 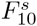 and 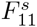 are two Euclidean distance-based features which capture the proportion of each telomere region in the combination to the perimeter of the rectangle made by connecting the 4 candidate points. During our investigation, we observed a significant improvement of the accuracy of classification by the inclusion of these two features.

Let *d*(*p; q*) denote the Euclidean distance between the points p and q. Similarly let *l* (*p; q*) represent the length of the curve between p and q, which are points from the set *D^P^*. Then, for each contour partitioning combination in *T^P^* given by 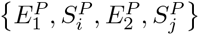 (where *i* and *j* are integer values such that 1 ≤ *i, j* ≤ 4 and *i* ≠ *j*), two main length measurement ratios (r1 and r2) are used for both calculating length based features, as well as for normalizing these features. 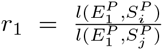 yields the chromosome width/length with respect to the anchor point 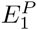 for the given contour partitioning combination (refer figure 3). Similarly 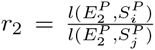 is calculated with respect to the anchor point 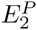. Then, the set of features *F^s^* for each contour partitioning combination is defined as follows,

1. 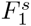 = 1 if the point 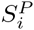 belongs to a skeletal end point (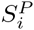 ∊ (*S* ∩ *C*)). Otherwise, 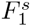 = 0.
2. 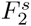 = 1 if the point 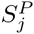 belongs to a skeletal end point (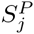 ∊ (*S* ∩ *C*)). Otherwise, 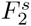 = 0.
3. 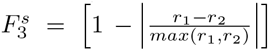 where 0 < 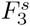 < 1. This calculates the chromosome width/length ratio for each anchor point and the difference between the two measures. Two similar fractions would result in a high value for the feature 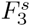.
4. 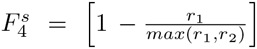 where 0 < 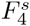 < 1. This calculates the chromosome width/length ratio with respect to the first anchor point (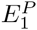). Except for smallest chromosomes at the highest degree of metaphase condensation, the telomere axis is shorter than the longitudinal dimension of the chromosome. Therefore, a lower length ratio measurement is a higher value for the feature 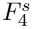 and is a desirable property.
5. 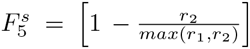 where 0 < 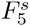 < 1. This is same as 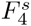, but from the other anchor point, 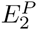.
6. 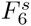 ratio of length of the chromosome to area of the chromosome. This provides a measure of elongation of a chromosome.
7. 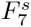 ratio value of perimeter of the chromosome to the area of the chromosome. This provides a measure of how noisy the object boundaries are.
8. 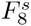 average of the curvature values *Φ_h_* of the candidates. The curvature is an important measurement of the saliency of the candidate points.
9. 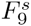: number of the negative curvature values (*Φ_h_* < 0) of the candidates points (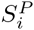 and 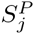). The telomere region end points are generally characterized by points with high convexity. The number of negative angles yield how concave the points of interest are.
10. 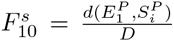 where 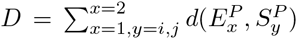. This feature calculates the normalized Euclidean distance between the anchor point 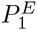 and the candidate 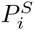 which makes up one telomere region.
11. 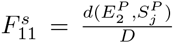 where 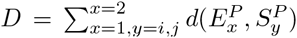. This is the same as feature 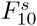, but calculated for the other anchor point.

A data set of 1400 chromosomes was collected from 40 metaphase cell images, which together yield 16,800 possible combinations of feature sets for contour partitioning. Three expert cytogeneticists marked the viable combinations of the salient points that capture the telomere regions for training the SVM classifier. The procedure involved training and testing with 2 fold cross validation (50% – train data, 50% – test data). We obtained values of accuracy, sensitivity and specificity values of 94%, 97% and 68%, respectively. The results demonstrate the ability of the feature set to effectively detect good combinations of candidate points for partitioning telomere regions. The slightly lower specificity suggests that some false positive telomeres were detection. However, this does not affect the accuracy of the contour partitioning, since the algorithm picks the optimal combination based on its rank rather than the classification label.

Correcting the deviation of the centerline for the effects of premature sister chromatid separation can be a difficult problem to solve. Once the best combination for the end points of the telomere region is selected, the telomere portions are segmented. Premature sister chromatid separation is detected from differences in the chromosome shape in the telomere region. This problem is solved with an algorithm that creates a set of features using functional approximation of the shape characteristics unique to premature sister chromatid separation and is derived from the coefficients calculated for each telomere [8]. A second SVM classifier is trained on these features to effectively detect these inherent shape variations of the sister chromatids. Once identified, correction is performed by extending the sample point (on the pruned centerline) to pass through the mid point of the partitioned telomere region. By getting the contour partitioned accurately, the correction process is significantly simplified.

### 2.3 Intensity integrated Laplacian method

The width profile, which is defined as the sequential width measurements along the centerline or the axis of symmetry of the chromosome, is an important measurement used for deriving the centromere location.

In previous studies, Laplacian-based thickness measurement algorithms have been utilized for cortical thickness measurement applications [18], [19]. This method takes the second order derivative of the object contour and solves the Laplacian heat equation to obtain the steady state condition. This is performed by splitting the object into two similar segments, maintaining them at different temperature levels and then allowing heat flow between the segments. The width profile of the chromosome is obtained by creating trace lines extending from one contour segment to the other following the static vector field created at steady state conditions on the potential field. The width profile calculated using this approach was observed to be more uniform and less noisy relative to analogous approaches based on centerline of the chromosome [20]. However, as a consequence of the sole dependency on the object contour, the width profile could still be adversely affected by irregularities in the object boundary.

We have previously presented an algorithm, in which intensity was introduced as an additional feature into the standard Laplacian-based approach to improve its accuracy by making it less dependent on the object contour [8]. The intensity information is included as an additional feature in the calculation in the form of a weighting scheme for the Laplacian kernel. This biases the flow of heat towards similar intensities. The intensity feature aids in minimizing the impact from irregular boundary of the chromosome segmentation by guiding the width profile trace lines to be contained within chromosome bands, which are regions with similar intensities.

### 2.4 Candidate point generation & metaphase centromere detection

In a previously described candidate-based approach, four candidate points were selected based on the minima values from the width profile [21]. However, this limits the number of possible locations that could be detected as the centromere location. Especially in cases where a high degree of sister chromatid separation is evident, limiting the search to just few candidates can have adverse effects. Therefore, we consider all possible local minima locations as candidates for the centromere location in a given chromosome, which are selected using the simple criteria given below.

Our notation *p* is used to refer to any other point(s), in general. Let the contour *C* be partitioned into two contour segments *C*^1^ (starting segment for tracing lines) and *C*^2^ (see figure 4). The width measurement of the normalized width profile at the discrete index λ (*W* (λ)) is obtained using the trace line which connects the contour points 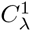 and 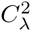 from the two contours *C*^1^ and *C*^2^. Then, the set of candidate points for the centromere location *p^C^* (which stores the indices λ), where the local minima conditions of *W*(λ – 1) < *W*(λ) < *W*(λ + 1) and *W*(λ – 2) < *W*(λ) < *W*(λ + 2) are fulfilled for all valid locations λ of the width profile. In cases where the above condition failed to secure any candidates (mainly on extremely short chromosomes), the global minima was selected as the only candidate. Next, the following two sets of indices are created to correspond with each given element *p^C^* (α) of *p^C^*,

**Fig. 4.**
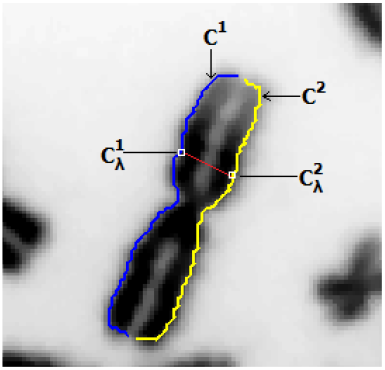
Illustrates an example where the contour *C* is split into two approximately symmetric segments *C*^1^ and *C*^1^. The width trace line, in red, connects the points 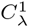 and 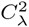 of the two contour segments.

– *p^mL^*(*α*) = *W*(*β*) where *W*(β) > *W*(γ), ∀γ < *p^C^*(α). Here *p^mL^*(*α*) stores the index of the global maxima for the portion (referred to as a regional maxima, henceforth) of the width profile prior to the candidate minima index *p^C^*(α).
– *p^mR^(α)* = *W*(*β*) where *W*(*β*) > *W*(γ), ∀γ > *p^C^(α)*. Similarly, *p^mR^(α)* stores the index of the global maxima for the portion of the width profile after the candidate minima index *p^C^(α)*.

Once the centromere candidate points *p^C^* and their corresponding maxima points *p^mL^* and *p^mR^* are calculated, the set of features *F^c^* are calculated as given below. A set of 11 features 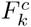 are proposed to train the third SVM classifier which will then be used to calculate the best candidate for a centromere location in a given chromosome. Features 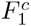 to 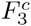 provide an insight on the significance of the candidate point with respect to the general width profile distribution. The normalized width profile value itself is embedded in features 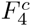 and 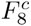 where the latter scales the minima based on the average value of the width profile. Features 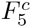 and 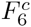 capture the contour curvature values that are intrinsic to the constriction at the centromere location. Features 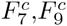 and 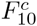 include distance measures which indicate the positioning of the candidate point with respect to the chromosome as well as to the width profile shape. Finally the feature 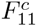 records the staining method used in the cell preparation. This gives the classifier a crucial piece of information that is then used to accommodate for specific shape features that may be the result of the particular laboratory procedure used to prepare and stain the sample.

Let *i* be a candidate member number assigned among the pool of centromere candidates. Also, let *d*(1, *i*) be the Euclidean distance along the midpoints of the width profile trace lines (centerline) from a telomere to the candidate point, and *L* be the total length of the chromosome. Then, the set of features *F^c^* are stated as below, where ∥.∥ yields the absolute value,

1. 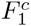∥*W*(*p^C^*(*i*)) – *W*(*p^mL^*(*i*))∥. This feature calculates the absolute width profile difference between the candidate and the regional maxima prior to the candidate point on the width profile.
2. 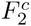∥*W*(*p^C^*(*i*)) – *W*(*p^mR^*(*i*))∥. This feature calculates the absolute width profile difference between the candidate and the regional maxima beyond the candidate point on the width profile.
3. 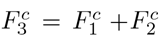 which calculates the combined width profile difference created by the candidate point.
4. 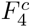 = W(p^C^(i)). This captures the value of the width profile (0 ≤ 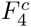 ≤ 1) at the candidate point location.
5. 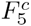 is the local curvature value at the contour point 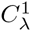 which corresponds to the current centromere candidate location (where λ = *p^C^*(*i*))
6. 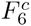 is the local curvature value at the contour point 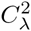 which corresponds to the current centromere candidate location (where λ = *p^C^*(*i*))
7. 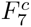 = *min* (*d*(1, *i*), *L* – *d*(1, *i*)) /*L*. Gives a measure where the candidate is located with respect to the chromosome as a fractional measure (0 ≥ 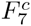 ≤ 0.5)
8. 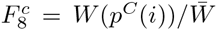, where 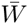 is the average of the width profile of the chromosome. This includes the significance of the candidate point minima with respect to the average width of the given chromosome.
9. 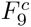 = *d*(*p^mL^*(*i*), *p^C^*(*i*))/*L*. This gives the distance between the candidate point location and the regional maxima value prior to the candidate point, normalized by the total length of the chromosome.
10. 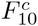 = *d*(*p^C^*(*i*), *p^mR^*(*i*))/*L*. This gives the distance between the candidate point location and the regional maxima value beyond the candidate point, normalized by the total length of the chromosome.
11. 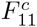 is a boolean feature used to indicate the staining process used during cell preparation. A logical ’0’ would indicate the use of DAPI chromosome staining while ’1’ would indicate a Geimsa-stained cell.

The detection of the centromere location assumes that each chromosome at least contains one centromere location within the chromosome. This is a reasonable assumption, since the centromere region is an integral part of chromosome anatomy which is normally retained in cell division, with the exception of acentric fragments produced by excessive radiation exposure, or rarely in congenital and neoplastic conditions. This assumption transforms the detection problem into a ranking problem in which we pick the best out of a pool of candidates. Therefore, this enables the same approach to be adopted that was utilized for the contour partitioning algorithm (section 2.2); ie. in which the distance from the separating hyperplane (*ρ*) represents a measure of goodness-of-fit for a given candidate. This metric reduces the multidimensional feature space to a single dimension, which inherently reduces the complexity of the ranking procedure for the candidate locations. Since the large margin binary classifier (SVM) orients the separating hyperplane in the feature space, the 1D distance metric directly relates to how well a given candidate fits into the general characteristics of a given class label. A detailed introduction to the candidate-based centromere confidence metric is provided in the following section.

#### 2.4.1 Candidate-based centromere confidence (CBCC)

Although existing measures of accuracy can establish performance of machine learning applications, these measures do not provide information on the reliability of the method. We developed a confidence metric for accurate detection of centromeres, which will be essential for assessment and ultimately adoption of this approach for diagnosis. We develop a Candidate Based Centromere Confidence metric (CBCC) to assess detection of a centromere location relative to alternatives. This value is obtained using the feature space derived via the classifier and the distance metric *ρ*. For a given set of candidate points, ie. centromeres, of a chromosome *p^C^*, the goodness of fit (GF) of the optimal candidate point 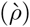 is obtained by calculating 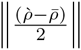, which is the average distance of all the remaining candidate points. In the ideal situation, the optimal candidate and the other candidates as support vectors for the classifier reside on opposite faces of the separating hyperplane (see figure 5). Therefore the optimal candidate distance 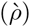 is ≈ 1, while the average of the remaining candidate distances 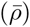 is ≈ –1. The GF value is truncated at unity, since exceeding this value does not add additional information to the metric.

**Fig. 5.**
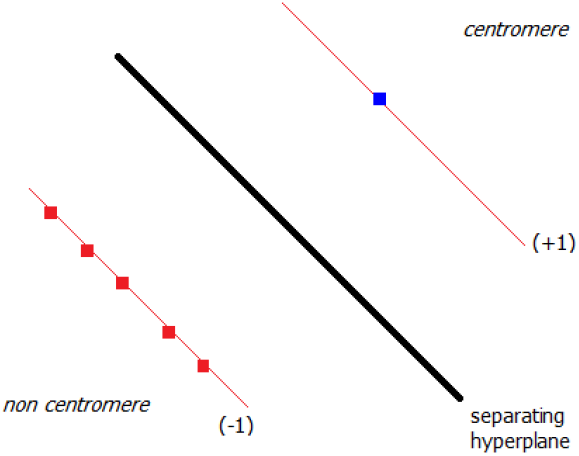
Shows the expected scenario for candidate-based centromere detection, in which 6 candidates are assessed by the SVM. The blue square represents the optimal candidate while the other five candidates are given by the red squares in the feature space.

## 3 Results

The complete data set used for developing and testing the algorithm discussed in this paper consists of 40 metaphase cell images, of which 38 consisted from irradiated samples obtained from cytogenetic biodosimetry laboratories and 2 were non-irradiated samples from clinical cytogenetic laboratories. The chromosome data set comprised images of 18 Giemsa-stained cells and 22 DAPI-stained cells. The cells with minimal touching and overlapping chromosomes (a good metaphase spread) was manually selected from a pool of 1068 cell images for this experiment. Then 40 cell images were selected to represent both DAPI (55%) and Giemsa (45%) staining methods. During ground truth evaluation, the expert was presented with the set of centromere candidates generated by the algorithm and was asked to select the candidate that closely represent the correct chromosomal location, while explicitly marking other candidates as non-centromeres. In cases where all the candidates suggested by the algorithm were incorrect, all the positions were designated as negative candidates. Intra-observer variability between experts (ground truth) was minimal, as the two laboratory directors differed in assessment in a single centromere out of > 500 chromosomes analyzed by both. The 1400 chromosome data set yielded 7058 centromere candidates. A randomly selected portion comprising 50% of this data set along with the corresponding ground truth centromere assignments were used for training a support vector machine for centromere localization. Next, the accuracy of centromere localization was calculated and is provided in Table 1. This provides a breakdown of the detection accuracy of the algorithm based on the presence or the absence of sister chromatid separation in the cell image for each staining method.

**Table 1.** The detection accuracy values for chromosomes used for the larger data set based on the staining method and the sister chromatid separation (SC Sep.)

| Chromosome morphology | Number of chromosomes | Number of accurate detections | Detection accuracy |
| --- | --- | --- | --- |
| DAPI without Sc Sep | 114 | 104 | 91.2% |
| DAPI with Sc Sep | 587 | 517 | 88.1% |
| Giemsa with Sc Sep | 699 | 599 | 85.6% |

Table 2 depicts CBCC values for accurately detected chromosomes as opposed to inaccurately detected chromosomes. It also includes a third category termed “All nonviable candidate chromosomes” (a subset of the inaccurate centromere detection category), where none of the candidates for a given chromosome were marked as capturing the true centromere of the chromosome. Table 2 Shows that CBCC metric demonstrates higher values in cases with accurate centromere detection.

**Table 2.** Shows that CBCC metric demonstrates higher values in cases with accurate centromere detection.

| Category | Chromo-<br>somes | Mean<br>($\mu$) | Std. Dev<br>($\sigma$) |
| --- | --- | --- | --- |
| Accurate detection | 1220 | 0.7861 | 0.3000 |
| Inaccurate detection | 180 | 0.3799 | 0.3293 |
| Nonviable candidates | 124 | 0.2696 | 0.2457 |

Figure 6 shows a representative sample of cases where the centromere was accurately localized. These cases include chromosomes with and without sister chromatid separation. The method does not detect centromere locations in all cases, some of which are impacted by the algorithm’s inability to fully correct for the adverse effects of sister chromatid separation (depicted in Figure 7).

**Fig. 6.**
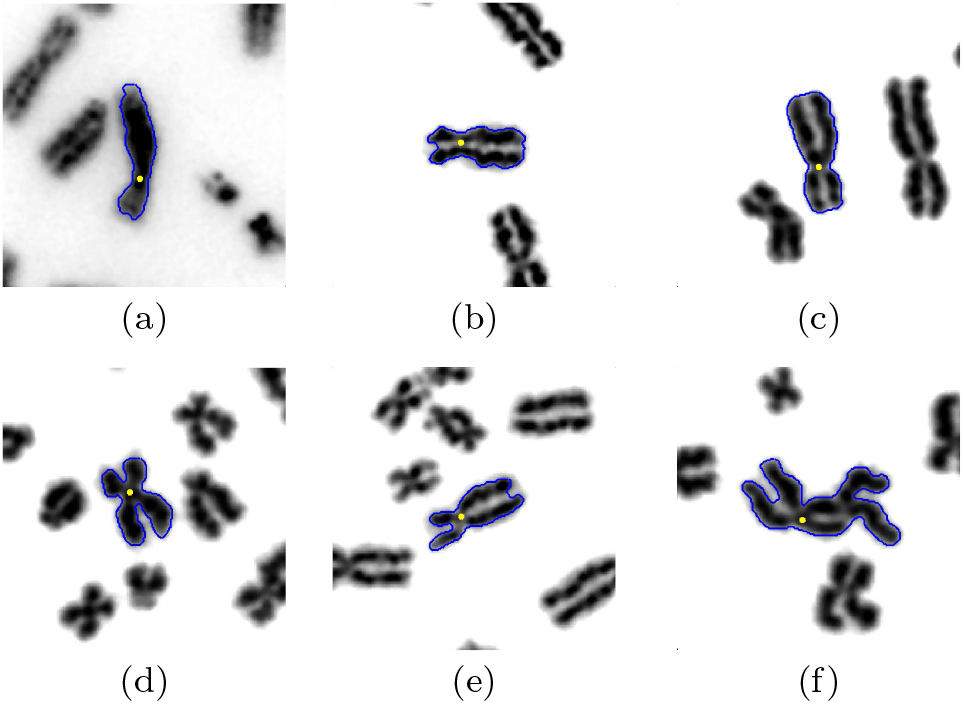
Demonstrates some sample results of the algorithm where the accurately detected centromere location (selected candidate) is depicted by a yellow dot while the segmented outline is drawn in blue. Figure 6(a) is a result of DAPI stained chromosomes while Figure 6(b)–(f) are results of Geimsa stained chromosomes. These results reported a CBCC measures of (a) 1.000, (b) 1.000, (c) 1.000, (d) 0.995, (e) 1.000, (f) 0.661 respectively.

**Fig. 7.**
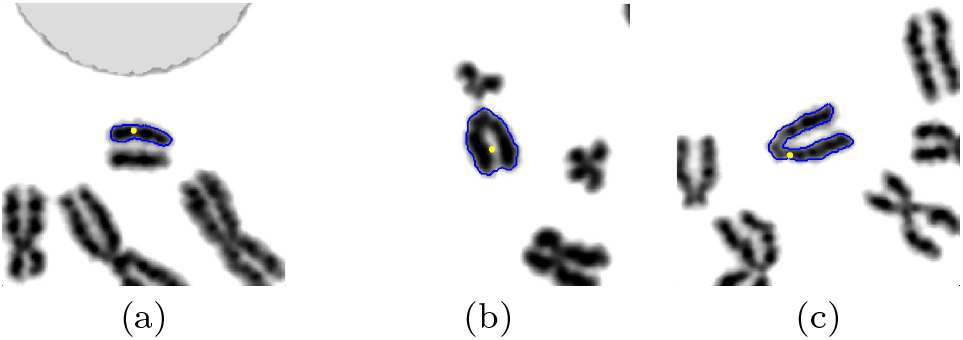
Demonstrates results where algorithm failed to yield an accurate centromere location. The detected centromere location (selected candidate) is depicted by a yellow dot while the segmented outline is drawn in blue. These results reported a CBCC measures of (a) 0.368, (b) 0.066, (c) 0.655 respectively.

## 4 Discussion

The candidate based approach for centromere detection used a trained SVM classifier based on half of the input chromosomes. The accuracy of the method was then tested using the remaining 50% of the data set (2 fold cross validation); accuracy, sensitivity and specificity were 92%, 96% and 72% respectively. Two fold cross validation was used instead of other methods such as the leave on out method, since it yields a reasonable estimation of the accuracy with a low computational cost. The higher sensitivity of this algorithm relative to our previous efforts [5] can be attributed to improvements in the performance of the classifier on both typical and sister chromatid separated chromosomes. The lower specificity is predominantly related to lower confidence detection by the integrated intensity Laplacian algorithm of centromeres in acrocentric chromosomes, in which the centromeric constriction is not readily apparent because of its close proximity to one of the telomeres.

The objective of this study was to accurately detect the preferred centromere location (points) for each chromosome, even though the SVM produces a set of candidate points that can each be classified separately. All candidates in each chromosome were analyzed separately and the best candidate from this set was selected based on the distance metric value (*ρ*) of which the results are produced in table 1. Upon testing, the algorithm accurately located a correct centromere location in 1220 of 1400 chromosomes (87%). It is notable that 124 of the 180 chromosomes that were missed were instances of non-viable candidate chromosomes. Some of these were caused by segmentation of acrocentric chromosomes, where the lighter intensity of the short-arm satellite regions were segmented out, while others were primarily the result of an extreme degree of sister chromatid separation, such that the pairs of telomeres from sister chromatids could not be unequivocally paired. The values in table 1 further suggest a slight reduction in accuracy for Giemsa-stained images, which contained significantly higher levels of sister chromatid separation and noisy chromosome boundaries.

The proposed method performed centromere localization accurately for chromosomes with high morpho-logical variations (see figure 6). From a machine learning point of view, figure 6(a), (b) and (c) are fairly straightforward centromere localizations. The CBCC values for all three cases were 1.000 which was truncated from an even higher value. This further validates the CBCC metric, indicating that the selected candidate is prefereable than the other candidates in the same chromosome. It is important to notice that the boundary conditions at the telomeric region of figure 6(c) is similar in appearance to those in figure 3 and figure 4. However, with further separation and intensity fading between the two sister chromatid arms, the segmentation algorithm could converge to a concave morphology in the telomere region that links the sister chromatids. Figure 6(e) represents such an instance where sister chromatid separation has had a significant effect on the chromosome segmentation. However, as a result of correcting for this effect, the algorithm has localized the centromere accurately with a CBCC value of 1.000. The chromosome segmentation in figure 6(d) demonstrates evidence of extensive sister chromatid separation and therefore the CBCC value is at 0.995, which still is a high value for the data set. The figure 6(f) represents a chromosome which is highly bent and also presents with very significant sister chromatid separation. Nevertheless, the algorithm was capable of localizing an accurate centromere location though the CBCC value was low (0.661), which indicates a less than ideal separation among the centromere candidates.

Some of the shortcomings of the proposed method are represented in figure 7. Most of these (86%) were observed to be cases where none of the candidates were deemed to contain the actual centromere. This was mainly due segmentation problems and very high levels of sister chromatid separation. Figure 7(b) depicts an example where the segmentation algorithm failed to capture the constriction in an acro-centric chromosome. The CBCC value in this example was as low as 0.066, which indicates that the algorithm selected a weak candidate for the centromere. Figure 7(a) demonstrates a case where extreme sister chromatid separation has caused the segmentation algorithm to treat each individual chromatid arm separately. This chromosome had a low CBCC value of 0.368, which is consistent with the acentric nature (morphological) of the separated arm. Figure 7(c) shows another impact of extreme sister chromatid separation, namely, the incorrect connection of the long arm of a pair of sister chromatids, leading to an apparent, bent chromosome, instead of detecting sister chromatid separation. The CBCC measure fails to distinguish this chromosome from a normal bent chromosome, but nevertheless yielded a relatively high value of 0.655.

Although it was not the focus of this study, we carried out a preliminary analysis of the capability of this algorithm to detect both chromosomes in a set of dicentric chromosomes, which were present among an excess of normal single centromere chromomes, due to irradation of some of the cytogenetic samples analyzed. The constriction at the second centromere is similar morphologically to the first centromere in these images, and therefore, it should be feasible that it be among the candidates found by the algorithm. We hypothesized that along with the optimal candidate, the second centromere was also expected to exhibit a short distance to the hyperplane and be well separated from the other candidates. These distances were compared for all centromere candidates, and probable dicentric chromosomes were identified by determining if the correct, ground truth centromeres were among the top four ranked candidates. The breakdown of the candidates which captured the second centromere location is given in table 3, where 20 cases (out of 31) reported the second centromere location as the second highest ranked candidate location. Among the 31 dicentric chromosomes present in the data set, the first candidate (the selected centromere) was accurate in all instances. There were only two instances where the second centromere was not among the top four candidates. In both of these cases, the chromosomes exhibited a high degree of sister chromatid separation. Nevertheless, the proposed method provides a good framework for detecting dicentric chromosomes in radiation biodosimetry applications.

**Table 3.** Shows that the proposed method ranked the second centromere in dicentric chromosomes higher in most of the cases.

| Rank of the second centromere | Number of cases |
| --- | --- |
| 02 | 20 |
| 03 | 6 |
| 04 | 3 |
| 05 | 1 |
| 06 | 1 |

## 5 Conclusions

We have described a novel candidate-based centromere detection algorithm for analysis of metaphase cells prepared by different culturing and staining methods. The method performed with an 87% accuracy level when tested with a data set of 1400 chromosomes from a composite set of metaphase images. The algorithm was capable of correcting for the artifact created by premature sister chromatid separation. The majority of chromosomes with centromere constrictions were detected with very high sensitivity. We have also tested an promising extension of the centromere detection algorithm to accurately identify dicentric chromosomes for cytogenetic biodosimetry. Loss of specificity in both mono and dicentric chromosomes was primarily the result of segmentation errors in acrocentric chromosomes, as well as in chromosomes with extreme degrees of sister chromatid separation. A better segmentation algorithm that addresses these challenging morphologies would further improve the detection accuracy of the proposed method. Furthermore, the telomere partitioning algorithm needs to be improved in order to handle chromosomes with extreme sister chromatid separation which are commonly encountered in radiation biodosimetry applications. In addition, an algorithm to accurately separate touching and overlapping chromosomes will also be required to fully automate this process. It is also necessary to analyze a larger data set to gauge performance of the proposed method.

The framework used for adding intensity into the Laplacian thickness measurement algorithm can be easily extended to include other features besides the calculation of chromosome width. Further investigation aimed at both improving centromere detection accuracy and applications of this algorithm to other detection problems is warranted. The Candidate Based Centromere Confidence (CBCC) was introduced as a measure for confidence in each centromere detection. However, this metric can be applied to any problem which required a selection of a candidate from a pool of candidates. We suggest that the CBCC metric may be extensible to indicate the relative quality of a given cell image or of a set of slide containing a set of metaphases cells from the same patient. If successful, the CBCC metric may eventually limit the amount of time required to evaluate samples both prior to and during centromere detection.

## Acknowledgements

Supported by the Western Innovation Fund (University of Western Ontario), Natural Sciences and Engineering Research Council of Canada and the DART-DOSE CMCR (5U01AI091173-02 from the US Public Health Service) and the Canada Research Chairs Secretariat.

---

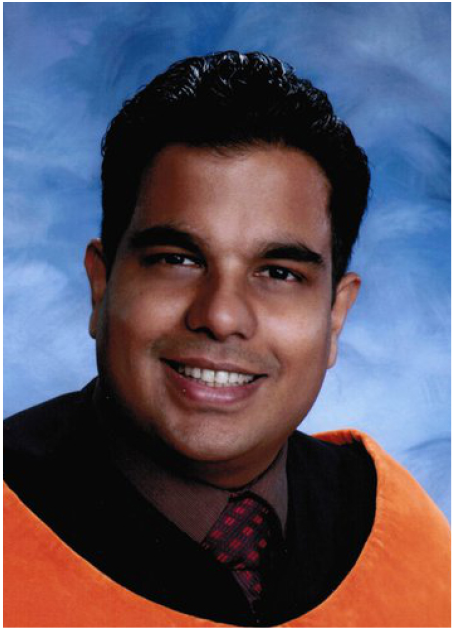

**Dr. Akila Subasinghe** (amsubasingheQsjp.ac.lk is a senior lecturer of Electrical and Electronics Engineering at the University of Sri Jayewardenepura, Nugegoda, Sri Lanka. He obtained a PhD of Electrical and Computer Engineering from the Western University, London, Ontario. He secured a BSc. Honors in Electrical Engineering from University of Moratuwa Sri Lanka and then a MESc. Electrical and Computer Engineering from Western University. Dr. Akila’s research areas include biomedical image analysis, biomedical instrumentation and machine learning.

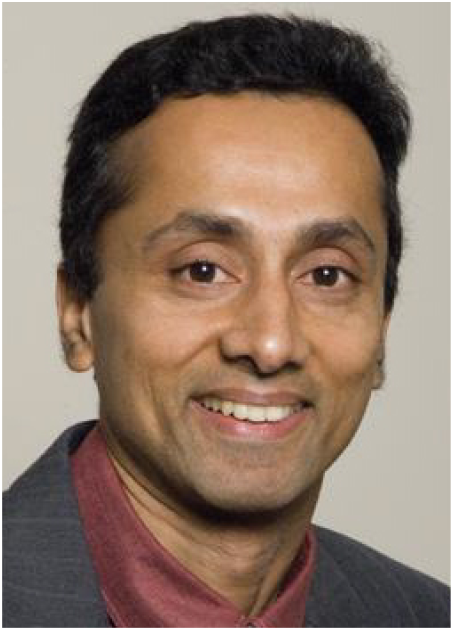

**Dr. Jagath Samarabandu**, is a professor of Electrical and Computer Engineering at the Western University, London Ontario. He is a Fulbright Scholar and obtained his MS and Ph.D. at SUNY Buffalo. His research areas include computer vision, pattern recognition, image understanding, machine learning and cyber-security. He has been with the university since 2000. Dr. Samarabandu’s research activities include image analysis, computer vision and pattern recognition. He has more than 20 years of academic and industrial experience in this domain.

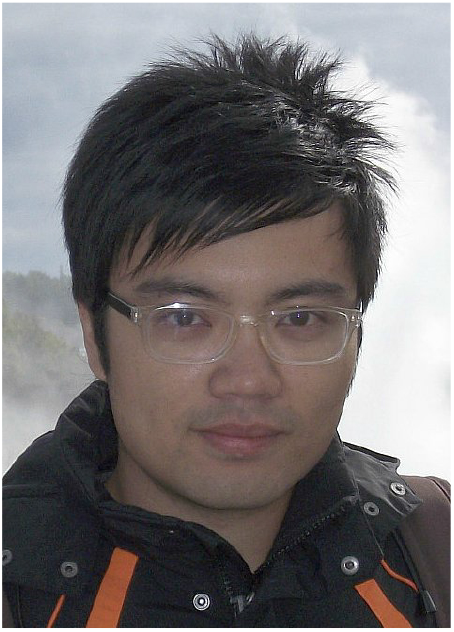

**Mr. Yanxin Li** is a software developer of Biochemistry Department at Western University, London, Ontario. He obtained his B.Eng in computer science and engineering at Zhejiang University and MSc in computer science at Western University. His research interests include parallel computing, machine learning and image processing. Hi is currently working on a research project to automate radiation dosimetry employing high performance computing.

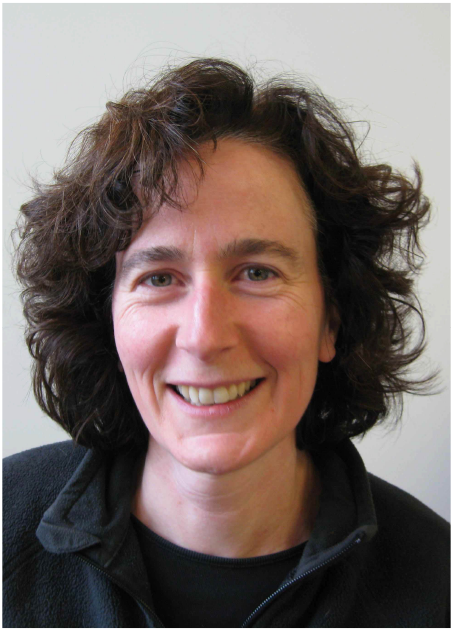

**Dr. Ruth Wilkins** earned a Doctor of Philosophy in Medical Physics from the Carleton University, Ottawa, Canada after which she joined the Consumer and Clinical Radiation Protection Bureau, Health Canada, as a Research Scientist. Since 2006, she has been the Division Chief of the Radiobiology Division at Health Canada. Her research involves the biological effects ionizing radiation in mammalian systems both in vitro and in vivo with a strong focus on cytogenetic biological dosimetry. She is currently the lead of the development of the Canadian National Biological Dosimetry Response Plan for large scale exposures to ionizing radiation.

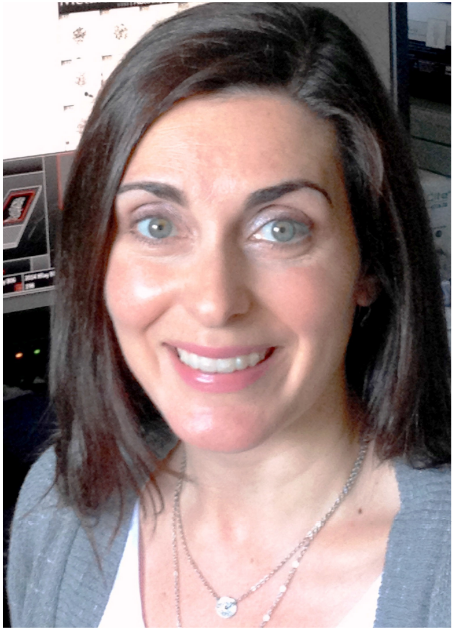

**Ms. Farrah Flegal** is a Research Scientist and lead of the Biodosimetry Program at Canadian Nuclear Laboratories (CNL). She is certified in clinical cytogenetics and molecular genetics by the Canadian Society for Medical Laboratory Science, the College of Medical Laboratory Technologists of Ontario, graduate of the University of Guelph with a BSc in Biological Sciences and from The Michener Institute of Applied Health Sciences with a DipHSc (Genetics). Her work at CNL involves being a partner in the Canadian Cytogenetic Emergency Network for emergency biological dosimetry as well as in research into the development of high throughput dose estimation methods for large scale exposures to ionizing radiation.

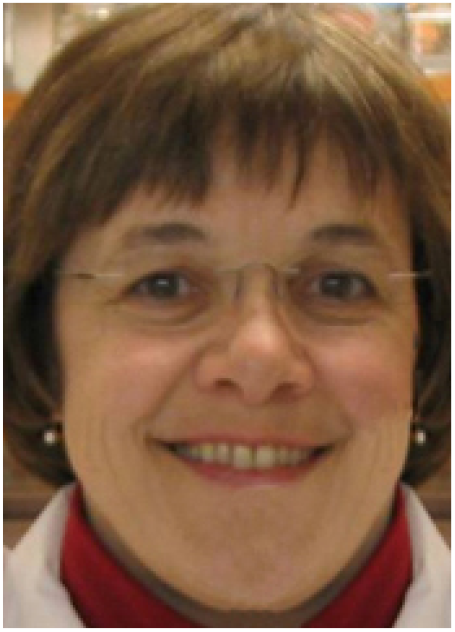

**Dr. Joan Knoll** is a professor at WU and Leader of Molecular Diagnostics Division, London Health Sciences Centre, Ontario (Ph.D.,, University of Manitoba; postdoc: Harvard University) and Chief Scientific Officer of Cytognomix. Dr. Knoll is board certified in cytogenetics and molecular genetics in the US and Canada. Dr. Knoll has more than 30 years experience in Cytogenetics including more than 20 years directing CLIA-certified and CAP or OLA accredited clinical cytogenetical laboratories. Knoll and Rogan founded Cytognomix based on their patents of cytogenetic/genomic technologies.

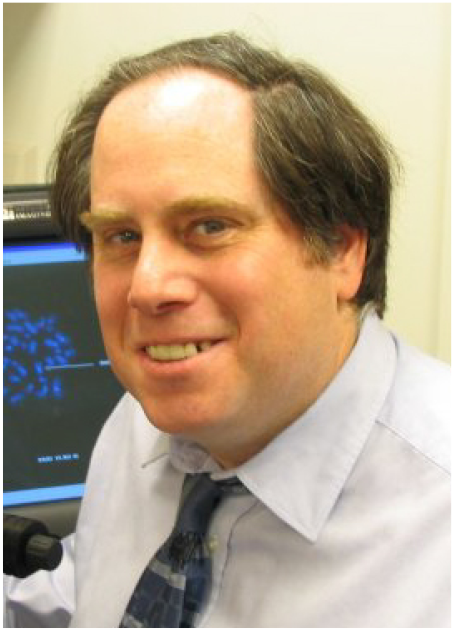

**Dr. Peter Rogan** Dr. Peter Rogan is Professor of Biochemistry, Computer Science, Oncology, and Biomedical Engineering at WU. He holds the Canada Research Chair in Genome Bioinformatics (B.S., Johns Hopkins; Ph.D., Yale University, postdoc: US National Cancer Institute), and is a prolific inventor and serial entrepreneur. He also cofounded Cytognomix Inc., which is developing software for automated detection of dicentric chromosomes in cytogenetic data. The company holds US Pat. No. 8,605,981 and other pending patents, which cover some methods described in this study. Rogan has more >20 years experience developing software applications that are widely used by molecular diagnostics laboratories.

## References

1. F. Cosoa, L. Vega, A. Becerra, R. Melndez, G. Corkidi, Medical & Biological Engineering & Computing 39(3), 391 (2001)

2. M. Munot, J. Mukherjee, M. Joshi, Medical & Biological Engineering & Computing 51(12), 1325 (2013)

3. M. Moradi et al., in 16th IEEE Symposium on Computer-Based Medical Systems (2003)

4. A. Vaurijoux et al., in Radiation research, vol. 178 (2012), vol. 178, pp. 357 – 364

5. A. Subasinghe A. et al., in International, Conference on Image Processing (ICIP) (2010)

6. R.M. Mohammad, Journal of Medical Signals and Sensors 02(02) (2012)

7. S. Jahani, S.K. Satarehdan, in International Journal of Biological Engineering, vol. 02 (1996), vol. 02, pp. 56-61

8. A. Subasinghe A. et al., in Biomedical, Engineering, IEEE Transactions on, vol. 60 (2013), vol. 60, pp. 2005 – 2013

9. T. Kobayashi et al., in IEEE Conference on Multimedia Imaging (2004)

10. P. Rogan et al., in Radiation Protection Dosimetry (2014)

11. M. Kass et al., International Journal of Computer Vision 1(4), 321 (1988)

12. C. Xu, J.L. Prince, IEEE Transaction on Image Processing 7(3) (1998)

13. L.J. Latecki, R. Lakamper, in DAGM-Symposium (1998), pp. 85-92

14. A. Subasinghe A. et al., in Seventh Canadian Conference on Computer and Robot Vision (CRV) (2010)

15. X. Bai et al., IEEE Transactions on Pattern Analysis and Machine Intelligence (PAMI) 29(03) (2007)

16. C. Xu, B. Kuipers, in Canadian Conference on Computer and Robot Vision (CRV) (2011)

17. L.J. Latecki, R. Lakämper, in Proceedings of the Second International Conference on Scale-Space Theories in Computer Vision (Springer-Verlag London, UK, 1999), pp. 398 – 409

18. M. Michael I. et al., NeuroImage 12(6), 676 (2000). DOI 10.1006/nimg.2000.0666

19. H. Haidar, J. Soul, NeuroImage 16, 146 (2006). DOI 10.1111/j.1552-6569.2006.00036.x

20. A. Subasinghe A. et al., in Electrical, Computer Engineering (CCECE), 2012 25th IEEE Canadian Conference on (2012)

21. R. Stanley et al., in Proceedings: a conference of the American Medical Informatics Association (1996), pp. 284-288

